# Sensing the Full Dynamics of the Human Hand with a Neural Interface and Deep Learning

**DOI:** 10.1101/2022.07.29.502064

**Authors:** Raul C. Sîmpetru, Andreas Arkudas, Dominik I. Braun, Marius Osswald, Daniela Souza de Oliveira, Bjoern Eskofier, Thomas M. Kinfe, Alessandro Del Vecchio

**Affiliations:** Department Artificial Intelligence in Biomedical Engineering, Friedrich-Alexander-Universität Erlangen-Nürnberg, Erlangen, 91052, Germany; Department of Plastic and Handsurgery, Friedrich-Alexander-Universität Erlangen-Nürnberg, Erlangen, 91054, Germany; Division of Functional Neurosurgery and Stereotaxy, Friedrich-Alexander-Universität Erlangen-Nürnberg, Erlangen, 91054, Germany

**Keywords:** Electromyography, Pose Estimation, Force Estimation, Neural Control of Movement

## Abstract

Theories about the neural control of movement are largely based on movement-sensing devices that capture the dynamics of predefined anatomical landmarks. However, neuromuscular interfaces such as surface electromyography (sEMG) can potentially overcome the limitations of these technologies by directly sensing the motor commands transmitted to the muscles. This allows for the continuous, real-time prediction of kinematics and kinetics without being limited by the biological and physical constraints that affect motion-based technologies. In this work, we present a deep learning method that can decode and map the electrophysiological activity of the forearm muscles into movements of the human hand. We recorded the kinematics and kinetics of the human hand during a wide range of grasping and individual digit movements covering more than 20 degrees of freedom of the hand at slow (0.5 Hz) and fast (1.5 Hz) movement speeds in healthy participants. The input of the model consists of three-hundred EMG sensors placed only on the extrinsic hand muscles. We demonstrate that our neural network can accurately predict the kinematics and contact forces of the hand even during unseen movements and with simulated real-time resolution. By examining the latent space of the network, we find evidence that it has learned the underlying anatomical and neural features of the sEMG that drive all hand motor behaviours.

## 1 Introduction

Information about the movement intent of a body part is encoded with the electrical signals generated by the action potentials that travel through the axons and the skeletal muscle fibers. During voluntary movements, the surface electromyogram (sEMG) [1] records the action potentials generated by current fields at the muscle fiber level after an activation signal sent by the final pathway of movement: the spinal *α*-motor neurons.

For kinematics acquisition, all notable methods are camera-based as they are intuitive and provide optimal results [2]–[5]. The main limitation of such systems is that they can only record movements after detecting changes in body position. Moreover, these systems restrict the subject’s movements so that they are within the camera’s field of view. In addition, when used with human subjects, special care must be taken to protect privacy by, for example, obscuring the face.

Therefore, when an animal or human interacts with an object or assumes a static posture, there is no information about the forces exerted by the biological system. If we want on-demand kinematics and kinetics recognition for humans, it becomes clear that we need to find something other than cameras for use at home or on the street, where we cannot set up a recording volume. There have been numerous relevant attempts trying to fill this specific need [6]– 37] by exploiting the EMG signals to reconstruct the movement intent, yet all have some limitations that hinder the access to the large number of degrees of freedom from the human hand. We will delve into the shortcomings of these works later in the discussion section, but for now, here is a brief overview of the limitations of current sEMG decoding systems: 1) errors that are too big for some or all of the fingers [13]– [16], [18], [20]– [23], [25], [26], [38], 2) achieving less than 14 degrees of freedom (DoF) that the hand needs for executing flexion and extension movements [14], [15], [19], [21]– [26], [28]– [30], [38], and 3) or not attempting unseen combinations of different movements [13]– [16], [18]– [26], [28], [29], [31], [38].

In this paper, we lay the foundation for robust kinematics and kinetics prediction by building a new machine learning model that is tailored for high-density surface EMG signals. We designed an experimental protocol that allows markerless acquisition of hand kinematics and high-density EMG systems during motion tasks covering virtually all functional degrees of freedom of the human hand (*>*20) during slow and fast movement speeds. This method allows continuous kinetics and kinematics prediction by recording only the myoeletrical activity generated by the extrinsic (forearm) hand muscles. Our model is supported by theoretical derivations and experimental results that demonstrate its effectiveness as a replacement for markerless machine vision in situations where it cannot be used, such as in outdoor environments. Further, it also has several relevant benefits that include the prediction of contact forces when there is no observable movement (e.g., holding a cup of coffee), and because of sensor-based nature it can be, in the future, embedded in a wearable garment in a minimally intrusive way. Because this method allows the prediction of the full hand kinematics, it is then possible to explore the neural code that governs the control of movements of the human hand as depicted by sEMG sensors. By studying the latent space of the neural network, we found that our model learned distinct anatomical features. We argue that the latent space constitutes a detailed sEMG representation of the neural code of movement impinging to the skeletal muscles. Our model was able to separate the individual hand digit movements and the synergistic activation (e.g., grasping and two-finger pinch) in a high-dimensional space. This work has important implications not only for human-machine interfaces, but also for the study of animal behavior, because in the future we will be able to implant electrodes under the skin and observe animal behavior and flexibility of control of multiple muscles in the wild. To provide a better overview, we have created a showcase website for our paper at this **[link](https://raulsimpetru.github.io/Sensing-the-Full-Dynamics-of-the-Human-Hand-with-a-Neural-Interface-and-Deep-Learning/)**.

## 2 Results

We acquired sEMG data from 320 electrodes placed on the extrinsic hand muscles together with 5 cameras simultaneously recording different hand movements at 120 fps from 13 human subjects. We recorded twenty different types of voluntary movements of the hand including individual digit flexion and extensions at two movement speeds (0.5 and 1.5 Hz), grasping, two- and three-finger pinches, and rotation of the fingers. Also, we recorded a “random movement task”, where subjects were instructed to move the hand freely (Video 1, Fig. 1). The instruction of the videos was performed with a virtual hand that was synchronized with the sEMG and the digital cameras. The videos were processed by DeepLabCut [3] and Anipose [39] to create 3D coordinates for each hand joint (Fig. 1B and C). From the 3D coordinates, we further calculated angles as a different mean of encoding the kinematics. For the kinetics experiments, we recorded the sEMG signals synchronously with the force measured by a dynamometer (COR2, OT Bioelettronica, Turin, Italy). The train/test splitting of the data is done differently for kinematics and kinetics, but the way we use it after splitting remains the same. For training, the bins are shuffled randomly. The kinematics have two test sets. The first set, called “test” (Videos 1 and 2), consists of the middle 20% of each motion recording (Fig 1E) without the random one. In total, this set is 2 min 23 s long. The remaining 80% are used as a training set. The second set, called “random movement” (Videos 3 and 4), consists of the random movement task. We use this set to test whether an untrained combination of motions, as it occurs in real life, can be reliably predicted by our system. The training set consists of all other motion tasks. The kinetics tasks (2- and 3-finger pinching) consist of 4 force ramps. The test set consists of the last ramp of both tasks (2 ramps in total), while the training set is formed of the remaining ramps (6 ramps in total). A machine learning model is trained with the sEMG as the only input. The ground truth comes from the cameras or the force sensor. For each test subject, a model was trained on a machine with 8x RTX 3080, 2x Intel Xeon Gold 6226R @2.9 GHz and 384 GB RAM for 50 epochs, which meant about 4.5 hours of training each.

**Fig. 1:**
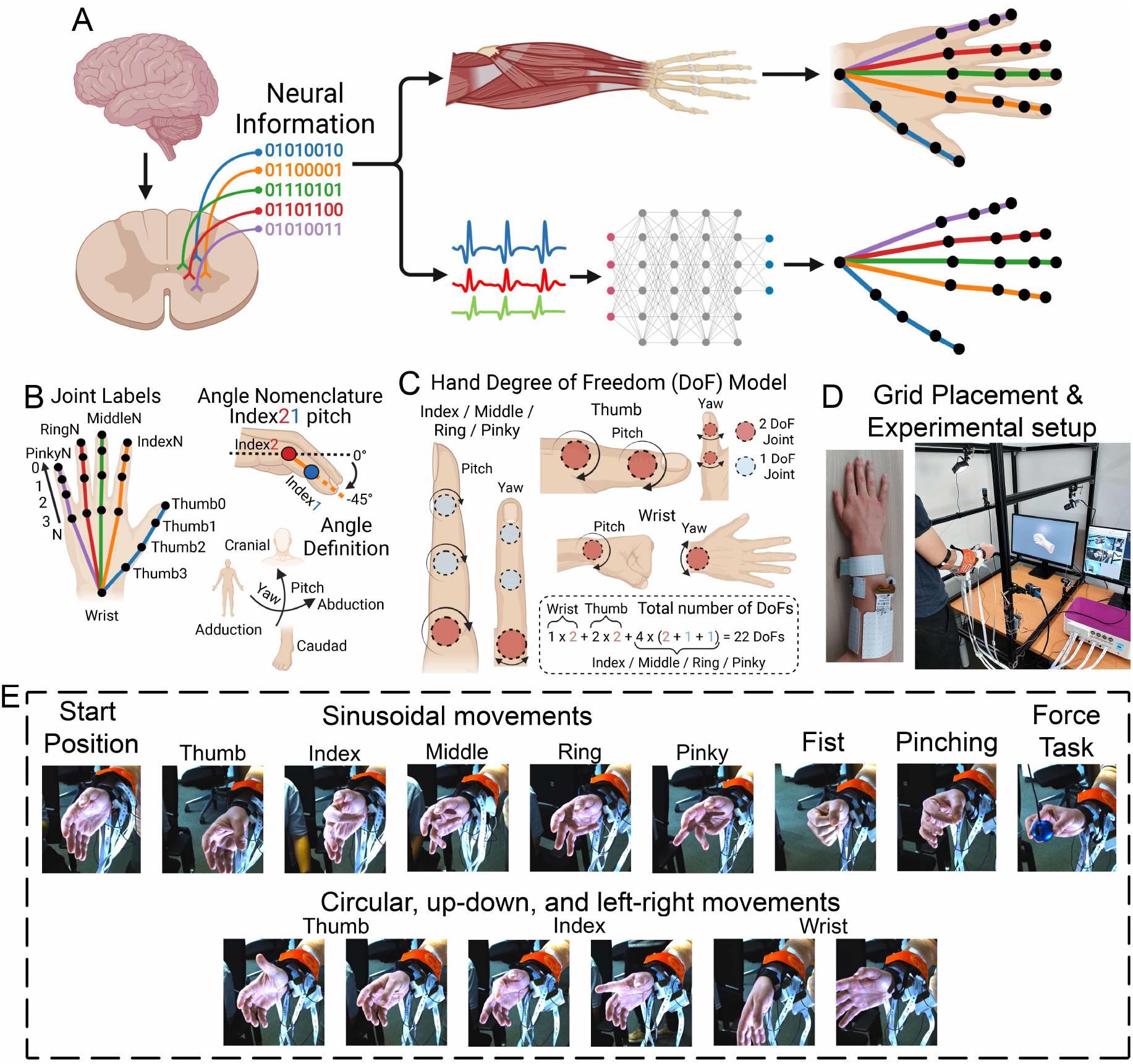
**A**. High-level overview of the study showing the neurophysiological pathway used for EMG generation together with the framework used for the training of the machine learning model. Hand movements are encoded by spinal and supraspinal pathways impinging on the final pathway of movement: the motor unit. The motor unit discharge timings encode the full dynamics of the hand that are ultimately amplified by the muscles and tendons. In our framework, the model learns the cumulative representations of the motor unit action potentials through highdensity grids of electrodes placed (320 electromyographic electrodes) on the forearm muscles. The deep learning model outputs the kinematics of the hand, therefore mirroring the musculoskeletal dynamics (i.e., the translation of the neural motor commands into movement and forces). **B**. Visual explanation of the joint labels and angle definitions. **C**. Hand degree of freedom (DoF) model. **D**. Pictures of the grid placement and the experimental setup. **E**. Representation of all executed movements.

### 2.1 Kinematics prediction

Although one type of kinematic encoding would have been sufficient, we chose to train our models using both the 3D spatial position of each joint and the joint angles as ground truth because we wanted to find out whether one or the other could generalize better on unseen data. Both output forms are filtered with a moving average filter of 150 ms.

#### 2.1.1 Three-dimensional hand estimation

Our AI model can output the entire hand stabilized by the wrist as 21 points in 3D space (Fig. 1B). The most intuitive error metric for spatial data is the mean Euclidean distance (MED). We used it to calculate the errors for both the test and random movement set, which are shown in violin plots (Fig. 2B-E). To compute the violins we take the mean across the joints, but not over time, and leave the wrist out of the calculation because it has a constant position due to the stabilization of the hand, which is not predicted and would therefore unfairly bias the result in a positive way.

**Fig. 2:**
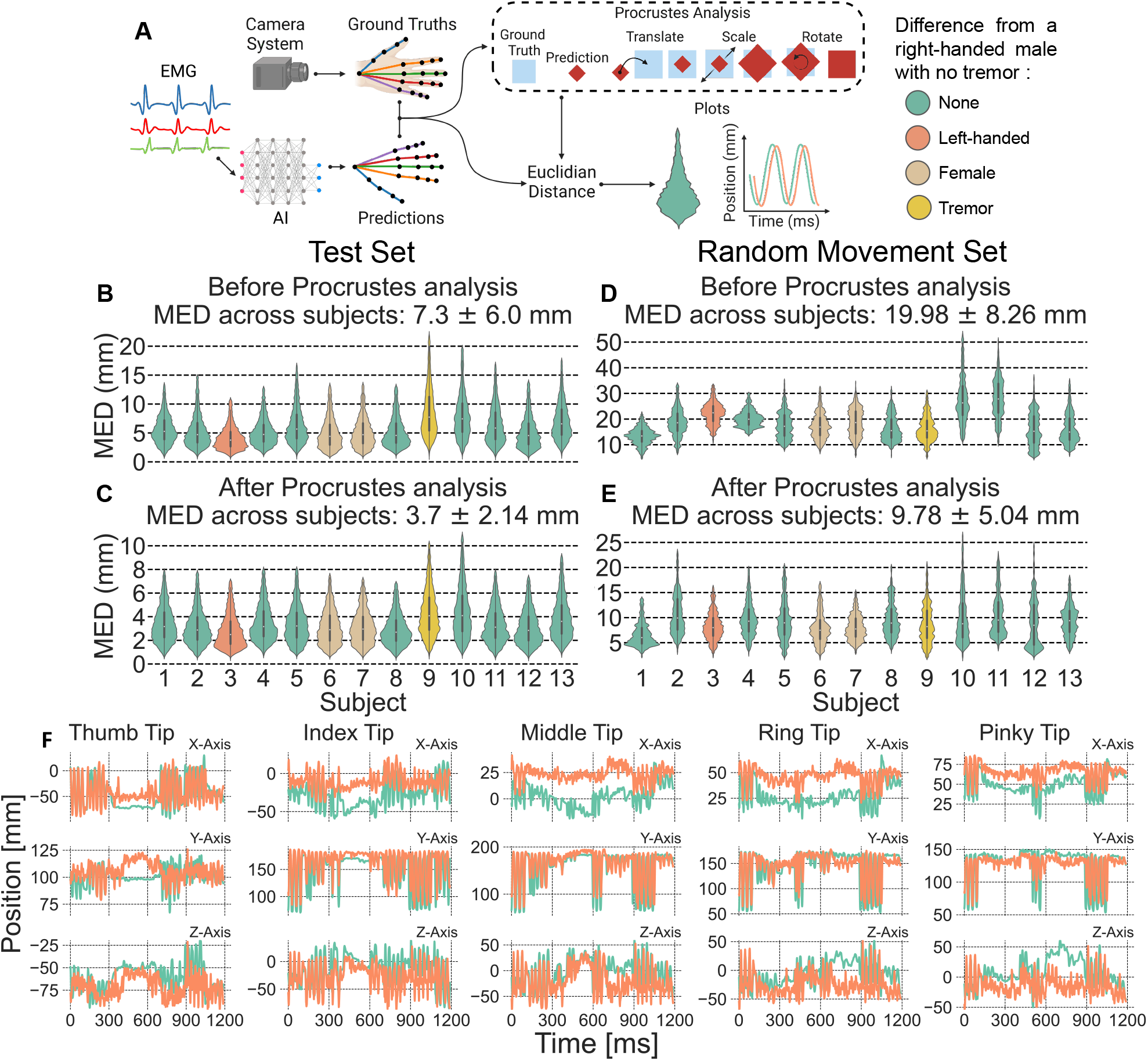
**A**. Graphical explanation of both the Procrustes analysis and the 3D points result plots. The violin plots show the distribution of all Euclidean distances during the entire duration of the set. This means that the Euclidean distances are averaged joint-wise for each time instance. **B. & C**. Violin plots showing mean Euclidean distance between the expected and predicted 3D joint coordinates both before and after the Procrustes analysis for the test set. **D. & E**. Same as B and C but for the random movement set. **F**. Comparison between expected and predicted positions of a few joints from subject 10 on the random movement set without Procrustes analysis. All joints can be found in Fig. A4.

We have noticed that although the model is outputting the correct movement trend, some small yet not insignificant jitter remains. To test this we used Procrustes analysis (Algorithm 1, see Methods section). Briefly, this method translates, scales, and rotates the output such that it may best fit the ground truth, while maintaining the same shape of the output and reducing the jitter. A graphical overview of both the Procrustes analysis and how we derive our result plots for the 3D outputs is shown in Fig. 2A. It is important to note that we use Procrustes analysis to show the prediction potential of the model if it was jitter-free, but we are not suggesting that this step could be replicated in real-time. Other filters that are better suited for real-time rendering can potentially achieve similar jitter reduction and should be tested in the future. However, it is important to note that the motion of the fingers can be clearly depicted even with the jitter.

Fig. 2B and C show the violin plots of the Euclidean distance between the predicted and expected 3D joint coordinates over the test set. The figures show the results before and after the Procrustes analysis. As mentioned, we also tested our model on a random motion task to evaluate if the system works in an environment more similar to a real life setting. Fig. 2D and E show the results of this analysis both before and after the Procrustes analysis. A few comparisons between predictions and ground truths from the random movement set of subject 10 can be seen in Fig. 2F, while in Supplementary Fig. A4 we have displayed all these comparisons.

We were able to achieve errors as small as 3.7 *±* 2.14 mm without jitter and 7.3 *±* 6.00 mm with the jitter across all 22 functional degrees of freedom of the hand. In the random movement task, the error increased to 9.78 *±* 5.04 mm without jitter and 19.98 *±* 8.26 mm with jitter (Fig. 2E), but this is to be expected since we did not train the neural network on this signal and the model is now going out of the distribution of the prediction values.

Although error metrics are important, they represent only a surrogate measure that is often difficult for humans to interpret and distinguish between excellent and only satisfactory results. Therefore we built a custom visual inspection tool using videos (example frame shown in Fig. 4) made from the original camera videos overlaid with the joint labels and 2 skeleton hands.

**Fig. 3:**
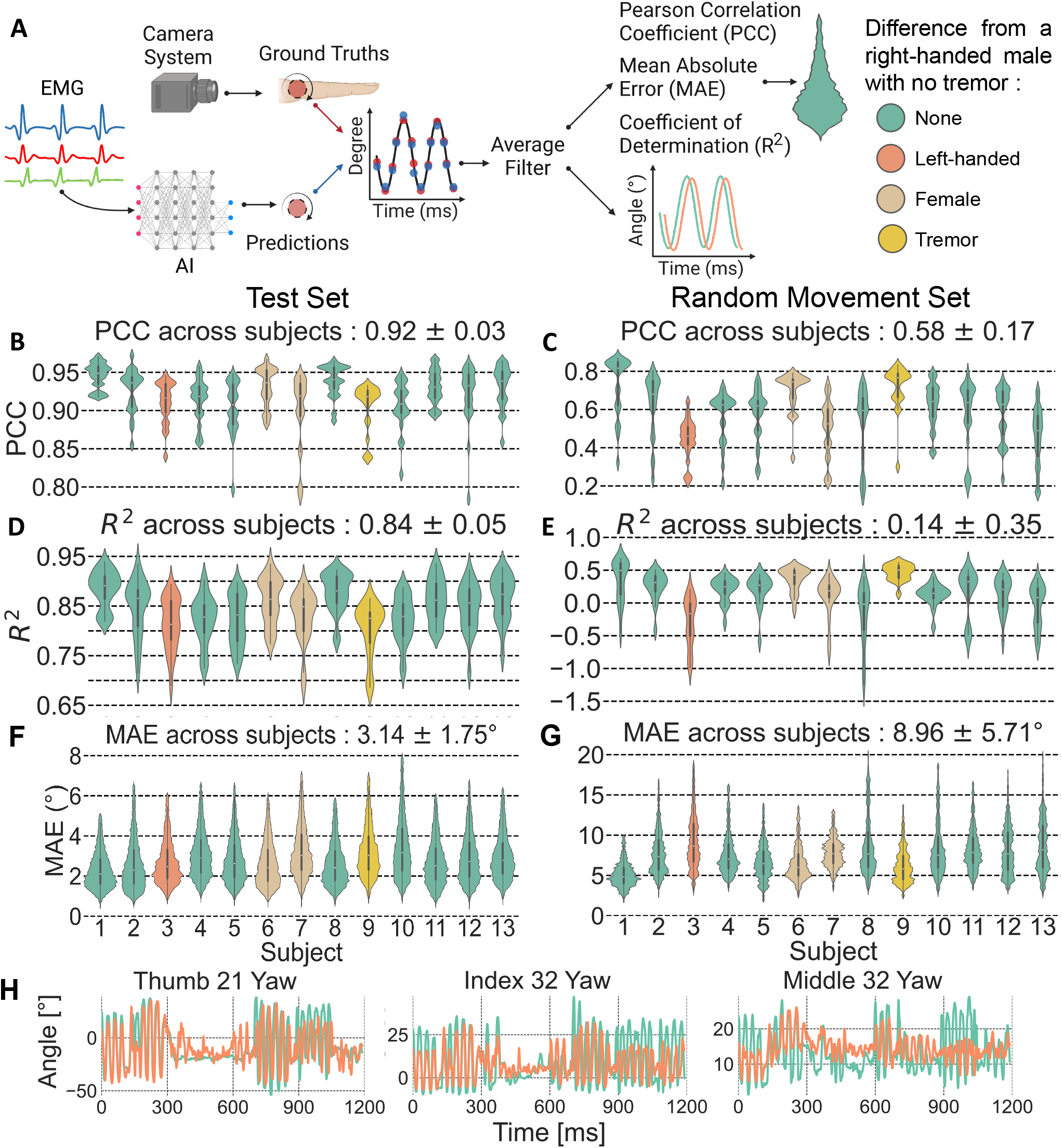
**A**. Graphical explanation of the angles result plots. **B. & C**. Violin plots showing the Pearson’s correlation coefficient (PCC) between the expected and predicted joint angles for the test and random movement set respectively. **D. & E**. Violin plots showing the coefficient of determination (*R*^2^) between the expected and predicted joint angles for the test and random movement set respectively. **F. & G**. Violin plots showing the mean absolute error (MAE) between the expected and predicted joint angles for the test and random movement set respectively. **H**. Comparison between expected and predicted angles of a few joints from subject 10 on the random movement. The angle names (Fig. 1B) are defined by 2 numbers. The first one is the origin and the second the joint to which the angle is calculated. All joints can be found in Fig. A5.

**Fig. 4:**
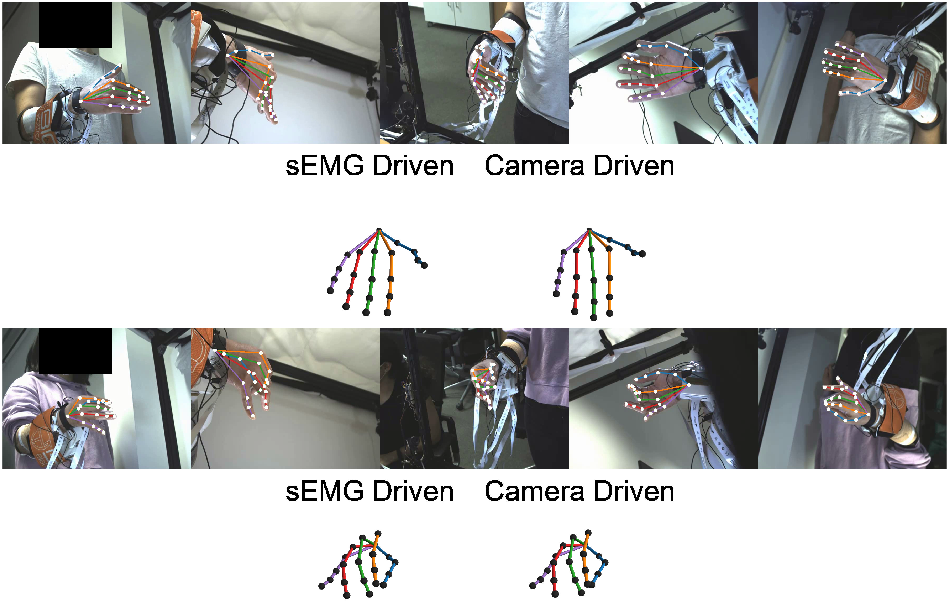
Example images from the videos used for visual inspection showing subject 1 flexing the wrist (top) and subject 7 performing a 3-finger pinch (bottom). The sEMG-driven hands use the 3D point output of our network, while the camera-driven ones use the 3D coordinates of Anipose [39]. All of our videos are accessible at this **link**.

The predicted hand is the 3D points output of our model, while the expected hand is determined by the coordinates obtained from the cameras. We provided the videos from subject 1 and 7 for both the test and random movement sets and made them available at this **link**. Also, we display the top 10 most active kernels and their activation maps across the EMG grids in Video 5 titled “Visualisation of the model kernel activities”.

To further explore the results, we developed an interactive online interface where the user can interact with a model trained on subject 1’s data to see the difference between the expected and predicted output. The model can be accessed at this **link**. We have then implemented the model in a real-time simulated scenario, by streaming the high-density sEMG in the same bins as processed in offline experiments. As in our previous conference paper [31] we found very similar results with this improved version of the neural network. We were able to consistently predict hand movements in simulated online settings every 31.25 seconds resulting in 32 predictions per seconds.

#### 2.1.2 Joint angles

For the angle outputs, we chose the Pearson correlation coefficient (PCC) (Fig. 3B and C), the coefficient of determination (*R*^2^) (Fig. 3D and E), and the mean absolute error (MAE) (Fig. 3F and G) as the error metrics for a robust overview of the predictions. Fig. 3A shows the overview of the analysis and prediction accuracy of the angle kinematics. Fig. 3F shows a few joint angles comparisons for subject 10 from the random movement set. The other joint angles can be seen in Fig. A5. The achieved mean PCCs are 0.92 *±* 0.03 and 0.58 *±* 0.17 for the test and random movement set respectively. The *R*^2^ for the test set is 0.58 *±* 0.17 and 0.14 *±* 0.35 for the random movement set. For the MEA we achieved 3.14 *±* 1.75° for the test and 8.96 5.71° for the random movement set. Similar as with the 3D points, all violin plots show the distribution over the duration of the set, with the data for each time point averaged over all angles.

### 2.2 Kinetics prediction

The force output is evaluated (Fig. 5A) exactly the same as the angles are - using PCC, *R*^2^, and MAE (Fig. 5B-D).

**Fig. 5:**
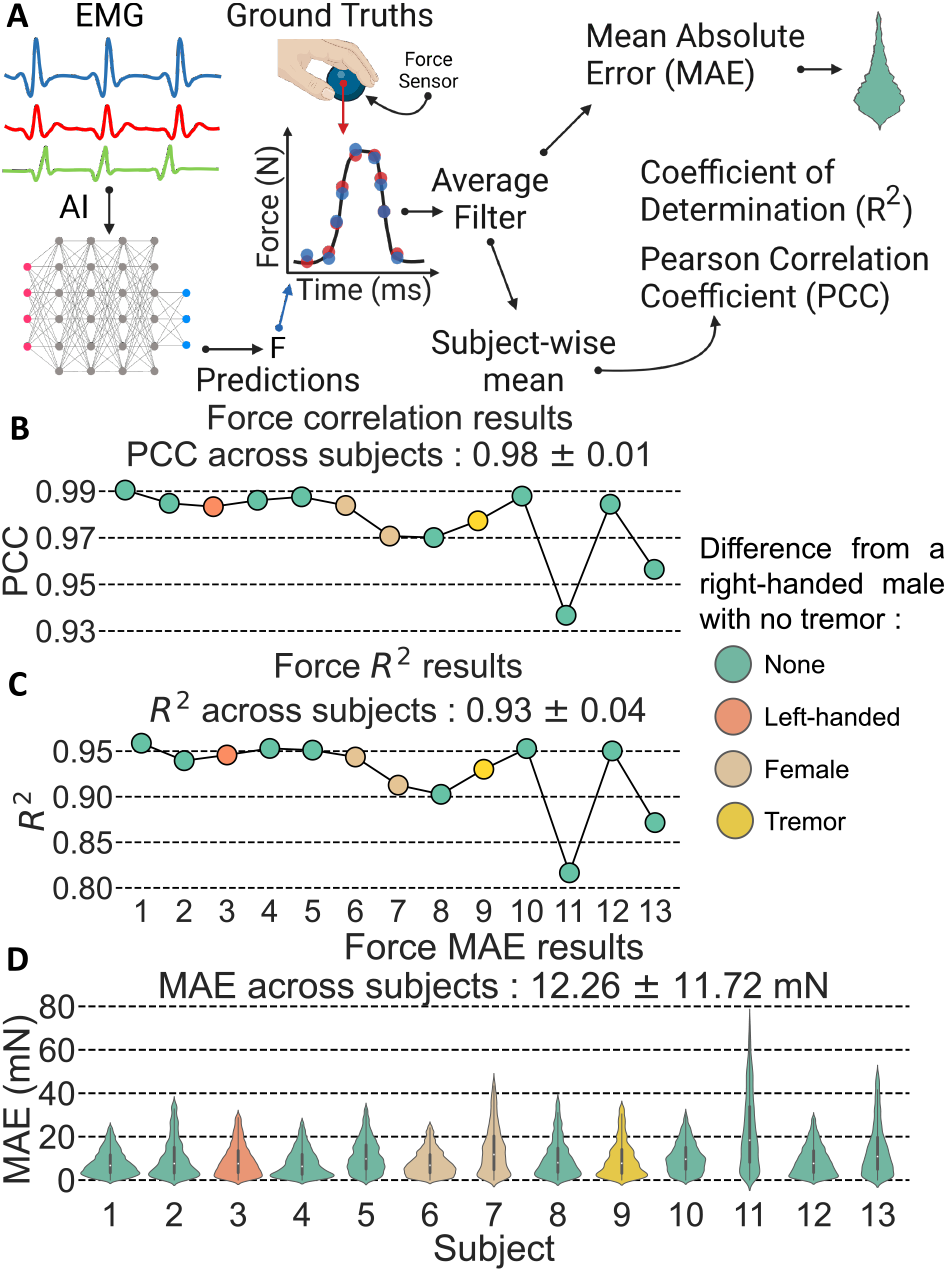
**A**. Graphical explanation of the force result plots calculated between expected and predicted force. **B**. Point plot showing the Pearson correlation coefficient (PCC). **C**. Point plot showing the coefficient of determination (*R*^2^). **D**. Violin plot showing the mean absolute error (MAE).

We recorded the force of the digits with an instrumented force sensor (Fig. 5A). The experiments included isometric ramp contractions in the full force range. As theoretically predicted, the neural network was able to predict the force output with an almost perfect correlation across all subjects (Fig. 5B-D).

### 2.3 EMG input comparison

We then looked at how different EMG input types affect the performance of our system for the 3D points output (Fig. 6). The input types are as follows:

**Fig. 6:**
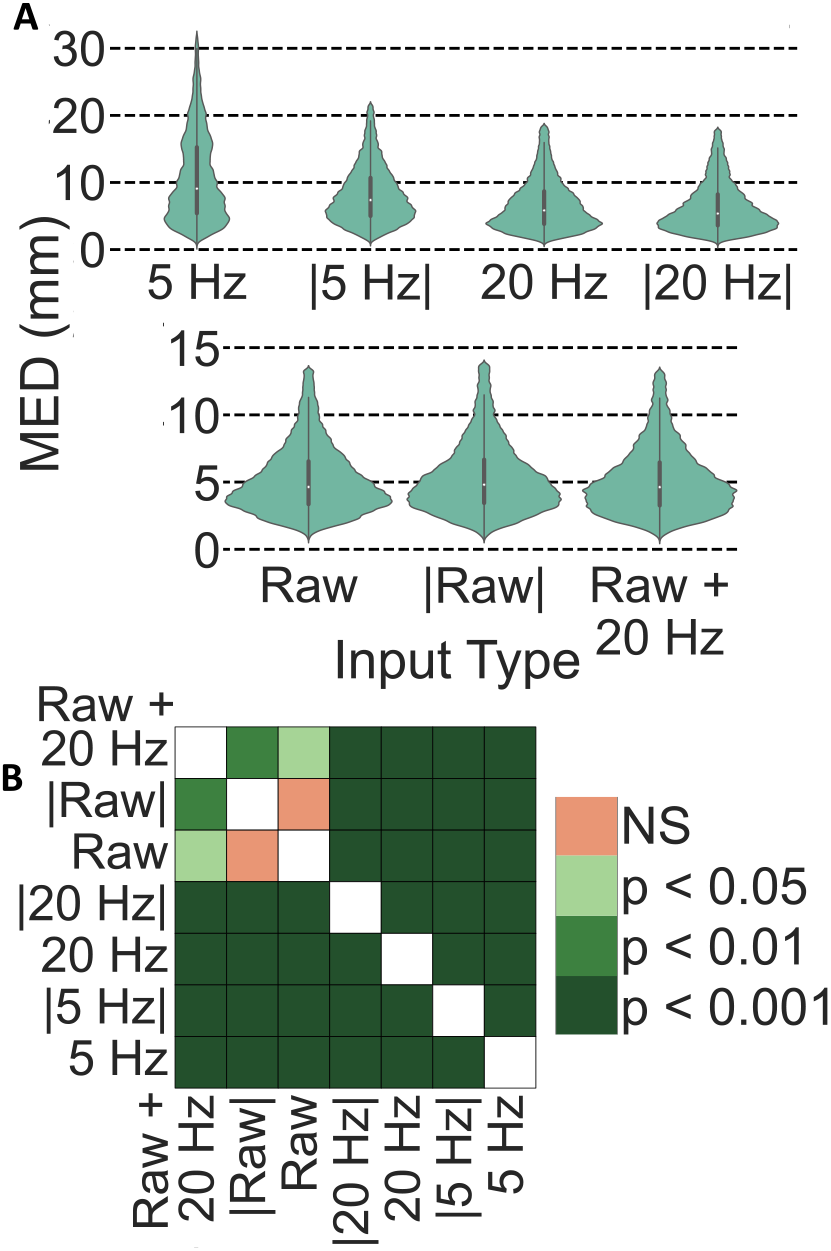
**A**. Comparison between different input types. The input types are described as : “x Hz” is the x Hz lowpassed filtered EMG, “Raw” is the unfiltered EMG, “| |” means that the EMG has been rectified, and “Raw + 20 Hz” is the input type we chose, which uses both the unfiltered and the 20-Hz low-passed variant of the same EMG. **B**. Statistical significance matrix calculated with paired t-tests with Bonferroni correction.

- 5 Hz - 5 Hz low-pass filtered EMG
- |5 Hz| - 5 Hz low-pass filtered rectified EMG
- 20 Hz - 20 Hz low-pass filtered EMG
- |20 Hz| - 20 Hz low-pass filtered rectified EMG
- Raw - unfiltered EMG
- |Raw| - unfiltered rectified EMG
- Raw + 20 Hz - unfiltered and 20 Hz low-passed filtered version of the same EMG

We have used a phase-free 4th order Butterworth filter to low-pass the EMG binwise backwards and forwards. The results shown are for the first 8 subjects only, as training models for the other subjects would unnecessarily increase the usage of our computing resources.

We display the mean Euclidean distance aggregated over all subjects as violin plots in Fig. 6A. To see if there is a statistical difference between them we ran paired t-tests with Bonferroni correction and display them as a heat map in Fig. 6B. As it could be theoretically anticipated (see Discussion section), the raw monopolar EMG signals were consistently the best input types for reconstructing all the kinematics of the human hand. This is not so surprising if we consider that the EMG bandwidth is in a relatively large range (*>*20 to *<*500Hz), and that sEMG motor unit action potentials in a muscle have high median frequencies (*>*40 and *<*300 Hz, [41]).

### 2.4 Mapping of the latent components learned by the neural network using densMAP

We investigated the latent space of our network with a non-linear mapping method. We projected the high dimensional latent vectors into 2 and 3D using densMAP [40]. Our model is made up out of a convolutional neural network (CNN) part whose output is then fed into a multi layer perceptron (MLP). Therefore, by looking at the output of the CNN we can understand the latent space.

DensMAP approximates high-dimensional manifolds and reduces them to human-understandable 2 or 3D visualizations. We chose densMAP because it does not assume that the data is uniformly distributed on a manifold and can therefore project the data more accurately as it would be in high-dimensional space. This is in contrast to UMAP [42], the algorithm densMAP is built upon. However in general the two algorithms work the same. Both UMAP and densMAP approximate the high-dimensional manifold that best fits the data, but UMAP directly searches for a lowdimensional projection, while densMAP first calculates the manifold densities in order to apply this information to the low-dimensional projection, preserving more of the original data structure. Further densMAP is capable of exploiting non-linear relations in the data, whereas methods like principal component analysis cannot.

A brief visual explanation of the analysis done can be seen in Fig. 7A. With densMAP we computed the 2D and 3D embeddings of each subject’s latent vectors (Fig. 7B) for the following movement tasks: flexing and extending each finger fast and slow, closing and opening the fist, and pinching with 2 or 3 fingers. The metric by which we separate the embeddings is the cosine similarity as this allows a detailed projection onto a sphere (3D) or circle (2D). The 3D embeddings of the subjects can be seen and manipulated at this **link**. We also manually (selection of hyperparameters for automatic detection of minima and maxima on the y-axis of task-relevant fingertips) labelled the flexion and extension parts of the kinematics so that we can display them in the clusters.

**Fig. 7:**
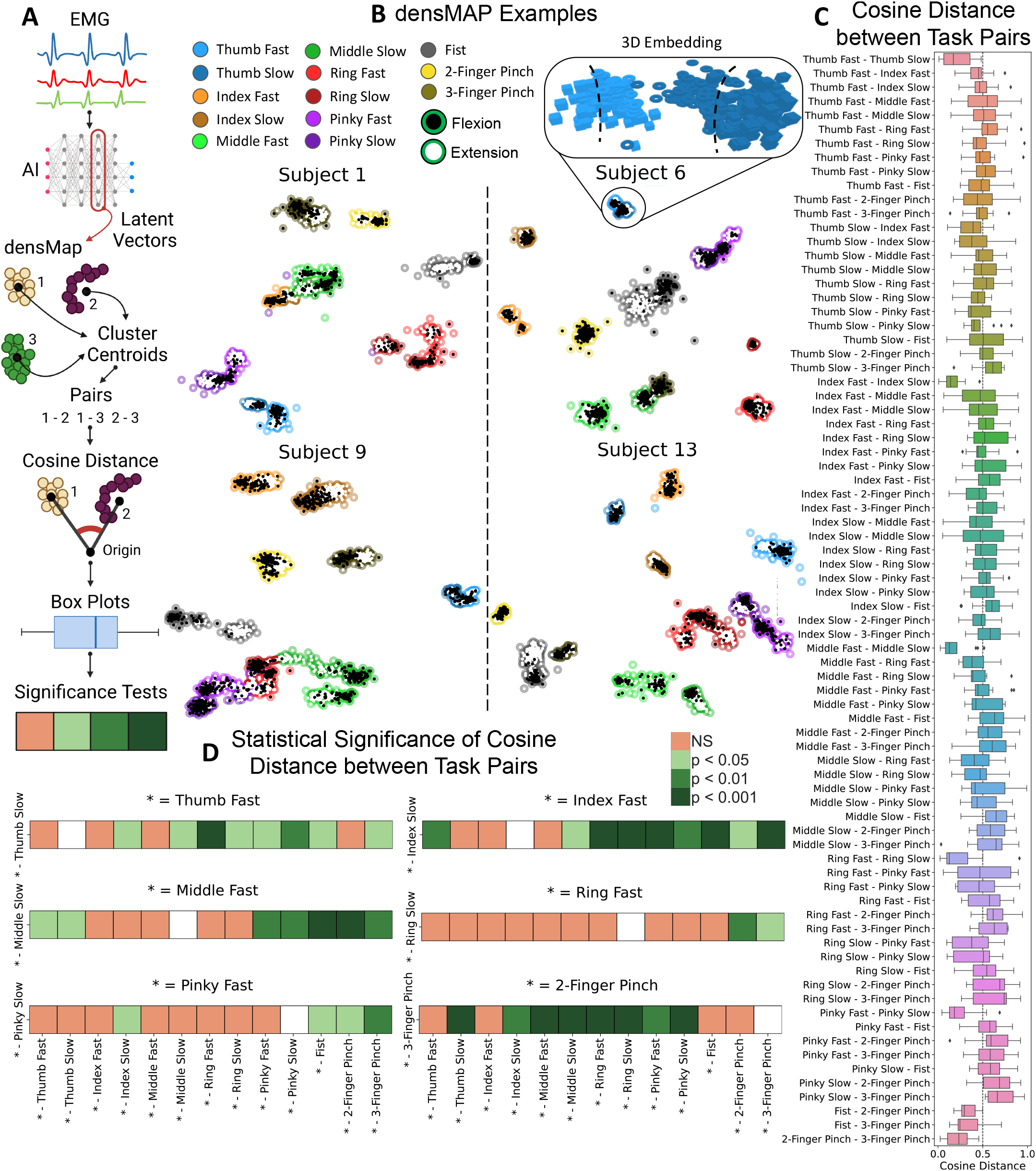
Analysis of the latent vectors using densMAP [40] to map them by cosine similarity. **A**. Simplified illustration on the analysis process. **B**. Example of subject embeddings in 2D. 3D versions can be accessed at this **link**. An example of a 3D embedding was shown to illustrate the fact that in the 2D projections it is sometimes difficult to distinguish the two clusters (most likely flexion and extension) that form a task, whereas in 3D they are more clearly visible. **C**. Box plots showing the cosine distance between all task pairs. **D**. Statistical significance was calculated using post-hoc pairwise t-tests with Bonferroni correction. Example interpretation: The cosine distance between “Ring Fast” and “Ring Slow” is significantly different from the pairs “Ring Fast - 2-Finger Pinch” and “Ring Fast - 3-Finger Pinch”.

From the 3D embeddings we compute the cluster centroids and calculate the cosine distance between each movement pair (Fig. 7C). Repetitions such as “Thumb Fast - Thumb Slow” and “Thumb Slow - Thumb Fast” have been removed as they both represent the same pair. We have to use the movement pairs because we cannot guarantee that densMAP projects the same movements in the same direction. However, we have assumed that the distances between the different tasks should be consistent across all subjects.

To then investigate whether the network has learned anatomical information about the hand we ran post-hoc pairwise t-tests with Bonferroni correction for different task pairs. We hypothesised that the distance between the fast and slow variants of flexing and extending each finger as well as between the 2 and 3 finger pinches should be significantly different from the rest of the pairs. The results of this analysis are displayed in Fig. 7D. An example interpretation of one of the results from Fig. 7D is that the distance between the task pair “Ring Fast - Ring Slow” is significantly different from the pairs “Ring Fast - 2-Finger Pinch” and “Ring Fast - 3-Finger Pinch”.

We also examined the difference between the cluster spread between fast and slow movements in finger flexion and extension. These results are displayed in Fig. 8 and discussed below.

**Fig. 8:**
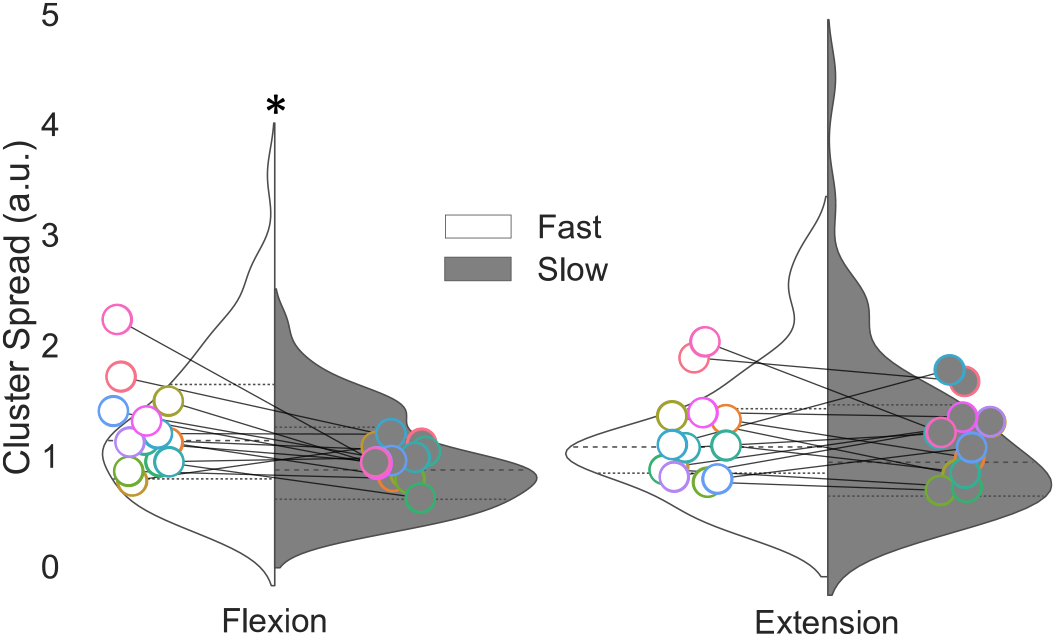
Cluster spread distance comparison between fast and slow finger flexion/extension movements. The subject means are overlaid on top. Significance has been checked with post-hoc t-tests after Bonferroni correction. \**p* = 0.0021

## 3 Discussion

In this study, we have collected and analyzed high-density sEMG signals during movements that covered virtually all the functional degrees of freedom of the hand at slow and fast movement speeds. We then developed and trained a neural network and demonstrated that the raw, unfiltered high-density monopolar sEMG contains all the information for recostructing the kinetics and kinematics of the human hand. Our results demonstrate that the model can accurately predict the kinematics on a test set and it is also able to predict movements on an unseen, random movement task. Additionally, our analysis of the model’s latent space using densMAP[40] reveals that the network has learned key aspects of hand anatomy and physiology. Finally, we support our choice of EMG input by testing various filtering combinations and evaluating their performance based on the resulting errors.

In this work, we aim to reliably decode the kinematics and kinetics of the human hand from sEMG. The estimation of joint forces (kinetics) from the sEMG signal has already been successfully addressed [43]–[46]. The sEMG is known to be highly correlated with the voluntary force level [47], [48]. The kinematics on the other hand is still an open question, which falls right into to prediction made by Hans Moravec [49] in what we today call Moravec’s paradox.

Moravec’s paradox stipulates that it is relatively easy to teach computer logic reasoning but hard to implement the perception and motor abilities of a one-year-old child. However, as described by the paradox, tasks that require complex dexterity and coordination, such as navigating and interacting with the world, can be challenging for computers to learn and perform. One way to teach computers how we move is to understand the neural signals that generate movement.

By recording the physiological signals associated to movement it may be possible to solve this paradox, and the literature is filled with examples [6]–[37] with some being more successful then others.

Regarding the prediction of movement from the sEMG, the current literature can be split into 3 distinct groups. The first and arguably most accurate (in terms of small errors and high prediction accuracy) group interprets the problem at hand as a classification task and wants to reliably distinguish between different hand postures. This approach is called gesture recognition [6]–[10], [17], [27], [32], [33], [35]–[37], [50]. Although achieving impressive results such as *≥* 90% accuracy on the Italian sign language [8] or 93.84% accuracy on 49 distinct poses [36], we believe that the method itself is lacking due to the users inability to chose the speed of executing the desired pose. The main problem with classification tasks is the lack of proportional control, e.g., it is not very useful to choose a specific grasp type to catch a ball but not being able to generate the required speed to catch it in time. Although multiple grab speeds could be selectable, there will always be an object that requires a class that is just between two existing ones to which the user does not have access to. This problem can only be solved by allowing the user to directly control the closing speed or by allowing so many speeds to be selected that the differences are effectively negligible. We consider the latter to be unfeasible, especially because the maximum of poses (as far as we are aware) that can be encoded with *≥* 90% accuracy is 65 [9] if we count isometric movements, or 52 [32] without isometric tasks.

Before investigating the other two groups that do allow the user to influence the speed of movement we need to understand something that is arbitrarily decided in almost all works - how do we encode kinematics in the first place? There are two ways of doing this in the literature: defining the joints of the hand as 3D points in Cartesian space [13] or calculating the angles between said joints [14]–[16], [18]–[26], [28]–[30], [38]. Just from the citations we listed it can be seen that there is an unbalance and clear preference for using the joint angles representation (Fig. 1C) over 3D points (Fig. 1B). It is obvious that encoding all degrees of freedom (DoF) of the hand as joint angles results in less variables as opposed to 3D points because if we have 21 joints to keep track of, they would result in 21 · 3 = 63 values while as joint angles we would get exactly as many values as DoFs (Fig. 1C). Since this small difference in output size should not be a real problem in modern systems, even when embedded, we decided to see if there is a difference in generalization on unseen data when we train our deep learning architecture with the two different encoding types. The two groups mentioned before are very similar as they both output the kinematic signal that the sEMG encoded and not predefined classes. Their main distinction is that one does it discreetly [11], [12] while the other continuously [14]–[16], [18]–[26], [28]–[30], [38]. From our review, it appears that the discrete version has not been explored much as it is subject to the same problem we found for gesture recognition. Even so outputting the discrete kinematic signal is a step forward in terms of user freedom and flexibility of contro, we can, for example, encode a 2D cursor that works almost flawlessly (*≥* 95% completion rate) with the flexion/extension of the wrist as left/right and opening/closing the fist as up/down [12].

Continuous prediction of the desired kinematics solves the problem by allowing the user to, theoretically, choose any speed at which to perform the movement. Regardless of the kinematics output format (3D points or angles) there are still a few problems with the solutions presented in the literature.

The first shortcoming that we use to immediately rule out a solution is the inability to decode enough DoFs. We believe that 14 DoFs are the minimum, as these are all the flexion/extension DoFs that the hand has and are sufficient to perform different types of grasping while still being useful enough even if fine-tuning through abduction and adduction is not given. Surprisingly a large number of previous works [14], [15], [19], [21]–[26], [28]–[30], [38] fall through because of this criterion and we are left with only a few contenders.

High errors on in-distribution tests are the next criterion for determining the accuracy of other methods. We decided to arbitrarily call averaged errors above 5° for joint angles and above 1.5 cm for 3D points as high by empirical experiments we performed on ourselves. These values are the smallest deviation that we believe could cause a substantial problem to the user. Further because a lot of works report the Pearson correlation coefficient, we select 85% as the cutoff because everything below would be rather counter intuitive for the user. For the same reason we select for the coefficient of determination (*R*^2^) 0.85 as the cutoff. Using this criterion we can further refine our selection by removing further works [13], [16], [18], [20]. We have also removed Quivira et al. [13] because although the reported errors seem to be below our threshold when considered as an average over all fingers, we can see that the thumb has high errors of about 2 cm.

We believe we have convincingly shown that a kinematic predictor is needed that not only achieves low errors, but is also capable of outputting all the DoFs of the hand, and that it must be tested for reliability and robustness on unseen data as well.

To fill the need for an improved kinematics predictor, we collected data from 320 sEMG channels together with the kinematics and kinetics of 13 subjects executing different hand movement tasks (Fig. 1E). Our deep learning framework was then able to estimate hand kinematics accurately (Fig. 2B-F, Fig. 4, and Videos 1-4) and achieved near-perfect force prediction (Fig. 5B-D).

Fig. 2B-E show the results for the 3D points output on the test and random movement sets respectively. Videos 1 and 2 show the results for the test set and Videos 3 and 4 for the random movement set for subjects 1 and 7.

Our model performs remarkably well on the test set showing results of 7.3 *±* 6.0 mm before and 3.7 *±* 2.14 mm after removing the jitter. We believe the noteworthy results are seen on the random movement task where our model achieves *≈*2 cm *±* 8.26 mm with jitter and *≈*1 cm *±* 5.04 when we remove it. These results together with the Videos 1-4 show us that the model is capable of predicting the kinematics reasonably well even on unseen movements without retraining.

When encoding the kinematics as joint angles we can see that the model predicts the test set in a relatively accurate way (Fig. 3B, D, and F). However, on the random movement set we see that the model performs poorly with an coefficient of determination (*R*^2^) of 0.14 *±* 0.35. From Fig. A2 we can see that the yaw angles (left-right) are the most difficult to predict. We suspect that it is much more difficult for the model to learn the biomechanical relationships from the joint angles because there, for example, flexion/extension has a largerrange of values than abduction/adduction, which could create confusion in the model. In view of these results, we recommend training the model to output 3D points and, if necessary, to calculate the angles in an additional step instead of having them outputted directly as they seem to be more stable on unseen data.

We have also tried to predict the kinetics using our architecture. It is well known that the EMG is highly correlated with the exerted joint force. In Fig. 5B-D we demonstrate that our model is capable of near perfect (0.98 *±* 0.01 correlation, 0.93 *±* 0.04 *R*^2^, and 12.26 *±* 11.72 mN of error) force prediction during 2- and 3-finger pinching, as anticipated.

To then see if our intuition about using the raw and 20 Hz low-pass filtered EMG as inputs holds any ground, we compared different input types and displayed them in Fig. 6. We conducted this test because we hypothesized that the common method in the literature of low-pass filtering the EMG (rectified or not) below 5 or 20 Hz [22], [24], [36], [51], [52] is an incorrect solution for a learning system like our deep learning model and for an accurate translation of EMG activity into kinematics. The result shows that the input type used in this study (raw and 20 Hz low-passed version of the same EMG bin) significantly outperforms all other types. This can be attributed to 2 factors. First, rectification removes important information that is picked up by the monopolar electrodes. Furthermore, prior rectification of the signal is not necessary for a learning model, as the mathematical operation that such a system should learn is trivial. We supplement the unfiltered (raw) EMG with the 20 Hz low-pass version because the muscle acts like a 5-6 Hz lowpass filter [53]. So by filtering the EMG for our deep learning model, we give some information about the effective neural drive that produces slow movements. From the result, we deduce that the combined input is most effective because the network likely learned the movement space (low-pass filtering) and the motor unit action potential overlaps used for specific movements (Video 5). It is likely that the neural network is learning ensembles of motor unit action potential overlaps, considering that the raw monopolar derivation shows the highest accuracy among all other inputs. This is due to the fact that motor unit action potential frequencies occur in a relatively large range (40-300 Hz) [41].

In the last result, we showed was the 2/3D projections of the latent vectors for all subjects using densMAP (Fig. 7). We conducted these experiments because we hypothesized that the model must contain some anatomical and/or neurophysiological information to achieve the predictions shown. Fig. 7D shows just exactly that. We can for example see that the cosine distance between the “Middle Fast - Middle Slow” movement pair is significantly different from all other “Middle Fast” pairs but the index and ring fingers. This is supported by a relatively similar anatomical position of the muscles and intrinsic synergistic nature. Another example would be the “Thumb Fast - Thumb Slow” pair whose distance is significantly different from every pair except the one containing the index and the 2-finger pinch. This is also because of the synergistic task during 2-finger pinch which includes movement of the thumb close to the index. Also, it is possible to note the similar embedding for the fist and 2/3-finger pinch actions. The speed of movement, in general, is also preserved but across digits, as it would be expected since the same muscles are activated during that task. Interestingly, the flexion and extension are however mapped in a very similar manifold with strong overlaps. We believe this is due to agonist/antagonist coupling (during the flexion and extension movements there is likely a high degree of common inputs to both agonist and antagonist muscles) that determines the manifold proximity. This analysis proves that the model has learned at least the anatomical information about the hand which likely constitutes a direct observation of the manifolds underlying theholdl code of movement at the sEMG level. We could use this in the future to monitor the rehabilitation of the hand by checking the embedding space for significant distances between the anatomically distinct clusters, as we believe that healthy hands should show the same significances that we have shown in Fig. 7D.

Furthermore, we show that related tasks are closer together, but also that flexion and extension are split into two clusters within the same movement task. We also demonstrate that the cluster spread is significantly different between fast and slow movements for finger flexion (Fig. 8), but not for extension. First, the spread must be different for flexion because when performing a fast movement, we need to recruit more motor units and thus have more statistical chances that all encode the same movement. Moreover, the available space for flexion is higher than extension due to hand anatomical constraints. The AI needs to fill more latent space for fast movements because there are more ways to execute them than for slow movements.

This shows that the model is learning the kinematical features from the sEMG in an unprecedented way with distinct features that were previously unknown.

## 4 Conclusion

In this paper, we present a new deep learning model capable of reliably extracting hand kinematics and kinetics from high-density sEMG data. The recording setup consists of 320 non-invasive electrodes spread over 5 64-channel grids (3 around the circumference of the forearm and 2 around the wrists) recorded synchronously with the hand kinematics by a high precision markerless camera setup and deep learning methods. Our results show that the model is even capable of acceptable predictions on all the functional degrees of freedom of the human hand on unseen data. Further, we also show that it has learned anatomical information about the hand by exploring and analyzing the latent space. Finally, we also test different input types for for the model and conclude that using the raw and 20 Hz low-pass of the EMG works best and achieves the smallest errors.

We contend that our model is implementable as a hand interface on already existing hardware and could be used for assistive device control and more immersive virtual reality controllers.

## 5 Methods

Before we get into the solutions required to archive our results, we would like to give the reader an extended high level overview to refer to if needed (Fig. A1).

Similar to our prior conference paper [31], we collected sEMG data from 320 electrodes and recorded simultaneously the hand movements using 5 cameras. We recorded 13 subjects out of which 11 were male and 2 female. Both female (6 and 7) and 9 males (subjects 1, 2, 4, 5, 8, 10, 11, 12, and 13) are right-handed. From the remaining 2 males, one (subject 3) is left-handed and the other (subject 9) is right-handed with essential tremor (neurological disorder that causes involuntary and rhythmic shaking).

The subjects were asked to perform 20 different movements (Fig. 1E gives a visual representation of these movements) for 40 s each:

- sinusoidal movement (flexion extension) of each digit at 0.5 and 1.5 Hz. These two frequencies have been experimentally chosen to represent slow/fast movements.
- closing and opening the fist.
- pinching between index and thumb.
- pinching between index, middle finger, and thumb.
- circular (clockwise and counterclockwise), left-right, and up-down movement of the index, thumb, wrist. We recorded 2 different sets of movement consisting of different sequences of the aforementioned movements.
- random movement (explain in detail below).

For the slow version of the movements we had *≈* 40 s · 0.5 Hz = 20 repetitions while for the fast version we had *≈*40 s · 1.5 Hz = 60 repetitions. The finger rotations, the 2- and 3-finger pinches and the opening and closing of the fist each had 40 repetitions. In total we had 760 movement repetitions or *≈*13 min 18 s of data per subject.

The random movement task that each subject is asked to perform, consists of a combination of previously done movements and was defined as follows:

- 6 times closing and opening the fist
- 6 times pinching with the index, middle and thumb
- 2 times flexing and extending the index finger
- 2 times flexing and extending the ring finger
- 2 times flexing and extending flexing the pinky finger
- 2 times flexing and extending flexing the ring finger
- 5 times pinching with the index and thumb
- 6 times closing and opening the fist
- rotating the index clockwise once

The movements were executed in the described order and took about 42 s to complete.

To aid and guide the subject in keeping a consistent movement frequency, a realistic virtual hand model created in Blender (Blender 3.0, Blender Foundation) was displayed on a monitor (Fig. 1C).

In addition to kinematics, kinetics synchronised with the sEMG during 2- and 3-finger pinching were recorded with a dynamometer (COR2, OT Bioelettronica, Turin, Italy). Before recording the force, we determined the maximum voluntary contraction (MVC) of each subject. We then recorded 4 force ramps for each type of pinch. One such ramp consists of 3 parts: the increase (from 0 to 40 % MVC - 10 s duration), the hold (constant 40 % MVC - 3 s duration), and the decrease (from 40 % MVC to 0 - 10 s duration).

The experiments were reviewed and approved by the ethics committee of the Friedrich-Alexander University (application no. 21-150 3-B) for compliance with the Declaration of Helsinki, and the subjects signed an informed consent form.

### 5.1 Data

#### 5.1.1 Acquisition

The sEMG is acquired from 3 electrode grids (8 rows x 8 columns, 10-mm interelectrode distance; OT Bioelettronica, Turin, Italy) placed around the thickest part of forearm and 2 around the wrist (13 rows x 5 columns, 8-mm interelectrode distance; OT Bioelettronica, Turin, Italy), proximal to the head of the ulna (Fig. 1C).

Prior to the electrode grids placement, the skin is shaved and cleaned with an alcoholic solution. The monopolar sEMG signals were recorded using a multichannel amplifier (EMG-Quattrocento, A/D converted to 16 bits; OT Bioelettronica, Turin, Italy), amplified (×150) and band-pass filtered (0.7–500 Hz) at source, prior to offline analysis. The signals were sampled at 2048 Hz and collected using the software OT BioLab (OT Bioelettronica).

While the sEMG signal is acquired, 5 720x540 pixel videos of the hand movement are recorded synchronously at 120 Hz by cameras (DFK-37BUX287, The Imaging Source™, Bremen, Germany) placed in different corners (upper/lower left/right) of a rectangular metal frame to capture multiple views of the same movement (Fig. 1D).

The videos were processed by a custom trained neural network using the DeepLabCut [3] framework. The network was trained on manually labelled frames of the hand and digit positions and outputs the 2D the hand joints. The 2D positions were then triangulated into 3D kinematics using the extension library Anipose [39].

The achieved mean absolute error (MAE) between user label position and predicted position is 3 mm, indicating high reliability in tracking of the hand digits kinematics.

From the 3D coordinates we then compute the yaw and pitch angles by defining one joint as the origin of a sphere coordinate system and the next joint in series as a point on said sphere, e.g. Index2 as origin and Index1 on the sphere (Fig. 1B).

To ensure accurate synchronization both inter cameras and intra cameras and sEMG recordings, a trigger signal was sent from a microcontroller (Arduino UNO, Arduino LLC) to each camera and the sEMG recording device.

The force is recorded by the dynamometer (COR2, OT Bioelettronica, Turin, Italy) at the same frequency as the sEMG (2048 Hz). To obtain the force information in newtons, we calibrated the sensor with several weights (10 g, 20 g, 50 g, 100 g, 200 g, 250 g, 500 g, 550 g, 700 g, and 750 g).

#### 5.1.2 Split

The kinematics data set was split in two ways to create two different sets of tests.

The first set, called “test”, is the classic approach where part of the data is used for testing and the rest is left for training. In our case, we took the middle 20% of each movement task (the random task was omitted) and used the remaining 80% for training. The test set is then *≈*2 min and 23 s long. This way we could check whether our model converges to a reasonable kinematic prediction or not.

To check if our model could perform in a test more akin to real-life we used the random movement task as the second test set and called it “random movement”. For training the models, we used all other movement tasks. It is important to note that we did not use transfer learning before attempting to predict the random movement. For clinical use, transfer learning would not be too much of a burden and could potentially improve the results significantly at the cost of a minor inconvenience in recording.

For the kinetics data we chose the last ramp from both the 2 and 3 finger pinches as the test sets. The remaining ramps were used for training.

### 5.2 Preprocessing

After the data has been recorded, we preprocess the raw dataset so that it can be used by our model. The preprocessing pipeline is shown in Fig. A3. The sEMG and videos are recorded while the hand moves according to a task. The videos and the EMG systems are synchronized by an analog input signal. The 3D points and angles were then oversampled from 120 to 2048 Hz to match the sEMG. The first 4.5 s of the recordings were removed, as this is the resting state used to check for EMG artifacts during recording. Since the termination of the sEMG recording is not automatic, the sEMG was cut to the length of the video. Then the recording was divided into bins of 192 samples in length (about 94 ms) with a shift between bins of 64 samples (about 31.3 ms). From this point on the kinematics and kinetics were treated differently from the sEMG.

#### 5.2.1 sEMG

The sEMG bins are copied, low-pass filtered forwards and backwards with a 4th order Butterworth filter, and depth stacked with the original unfiltered bins. The combined bins are then augmented with Gaussian noise and magnitude warping [54], providing a threefold data increase.

#### 5.2.2 Kinematics and Kinetics

We took the mean of the individual kinematics (3D points and joint angles) or kinetics (force) bins because they do not change significantly within 94 ms. This reduces the output form of our model from a matrix to only one vector per sEMG bin. The vectors are then normalised component-wise between -1 and 1 using the min-max normalisation function

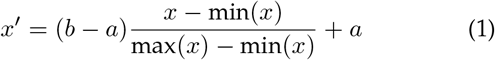

where a is the new desired minimum, b the new desired maximum, x the original signal and x^*′*^ the normalized signal. The minimum and maximum for the sEMG is only found within the bin, as otherwise the entire signal would have to be known, which would not be possible in a future real-time setup. For kinematics and kinetics, the minimum and maximum are determined from the entire signal. This function is necessary because differently scaled inputs can cause neural networks to become unbalanced and underperforming. Since the augmented sEMG bins must be paired with ground truth bins (kinematics or kinetics), we copied the original ground truth bins corresponding to the augmented bins.

### 5.3 Connection between EMG and cross-correlations

Before we go into the actual model architecture, we would like to set out the motivation for our choice by taking a closer look at the EMG generation theory.

#### 5.3.1 Modelling the EMG signal

The smallest unit that can be activated by the brain to perform a motor task is called motor unit (MU) [1], [55]. The MU consists of an *α* motoneuron and many muscle fibers innervated by its axon. One MU can innervate several muscle fibers, which means that the MU action potential (MUAP) does not have to be recorded at a specific point in the muscle, but can be recorded at several points simultaneously. Moreover, due to the properties of the volume conductor, the MUAP propagates through the muscle fibers and the biological tissue layers separating the action potential (source) and the recording electrodes, thus showing different waveform signatures in the high-density sEMG.

The EMG signal represents the sum of the individual MUAPs at different times [1], [48], [56]. These signals are recorded by non-invasive electrodes (usually arranged in a grid pattern) that are placed on the skin. The different firing patterns are used by the brain to encode information about the force required for a particular action. This process is called rate coding. In addition to the frequency of firing, the brain can also choose the number of motoneurons needed to perform an action, with more neurons meaning more power and fewer the exact opposite. This process is called MU recruitment and, together with rate encoding, they form the most important variables with which the brain can control muscle forces.

To model the described phenomenon we make use of following mathematical tools. We define the instances (also called spikes) *s*_*i*_ when a MUAP is observed by an electrode as a discrete Dirac delta impulse denoted as *δ*[ *t* ]. The sum of these spikes is then called the spike train *S*[ *t* ] of a MUAP. Since a MUAP can be observed for *I* different points in time *t*_*i*_, we define its spike train as

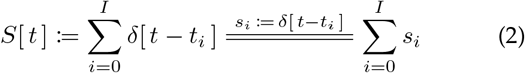

. To then produce a signal that has the MUAP at the points defined in its spike train, we can use the cross-correlation operation ⋆. Mathematically this operation is defined as

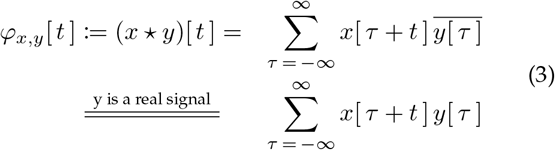

where *x*[ *t* ] and *y*[ *t* ] are 2 discrete signals and *φ*_*x,y*_[ *t* ] the cross-correlation between them. If *y*[ *t* ] is also a purely real signal with no imaginary component then it’s complex conjugate 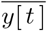 is the signal itself.

There are multiple ways of modeling the EMG, however most of them assume isometric movements [1] which are misleading if used in a dynamic context such as this paper wishes to address. Such inflexible models are usually used for decomposition and therefore assume that each MUAP must come from a different MU. To handle dynamic movements, we need to understand that the waveform of a MUAP can vary throughout the movement, even if it was generated by one and the same MU. These variations encode information about the location in the muscle, the orientation of the fibers, the speed of propagation of the action potential, the properties of the volume conductor as well as the properties of the recording system [1]. We call such variations MUAP observations and denote them with *h*[ *t* ].

Using all the information described so far, we can now start modeling the EMG signal one step at a time. A system with one electrode and one MU can be described by the following formula

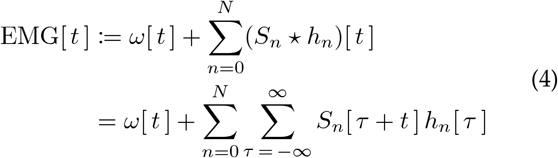

where *ω*[ *t* ] is the additive electrode noise, *S*_*n*_[ *t* ] and *h*_*n*_[ *t* ] are the n^th^ spike train (Eq. 2) and MUAP observation out of *N* variations of the MUAP of the single MU.

Although Eq. 4 is closer to anatomy we wish to reformulate it in a simplified yet mathematically equivalent way. Eq. 4 assumes that there are *N* different MUAP observations that each can have more then one spike where they are detected. We now constrain this by allowing each MUAP observation to have only one spike to which it corresponds, while extending the number of MUAP observations to *N*^*′*^ by allowing identical observations to count towards this length. Conceptually, this means that if for Eq. 4 we had only one observation detected at three different time points, in our new formulation we would have three observations, each corresponding to one of the different time instants. The reformulation then can be stated as

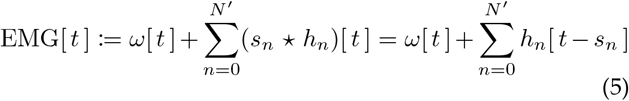

which essentially relieves us of one summation.

In order to extend our one-electrode system to *M* MUs, we have to redefine the observations and the spikes as follows

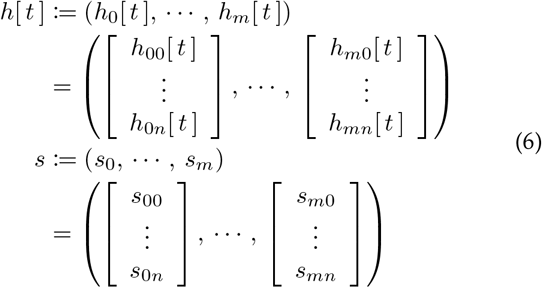

. The new observation and spike sequences each contain *M* elements for the *M* MUs. Each MU can have 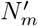 observations, which in turn means that each element of the sequence is a vector of that size. In this way, we can model each element of the sequence with different lengths, since some MUs can have more or less observations than others.

Using the new observation and spike sequences we can write the EMG signal for multiple MUs as

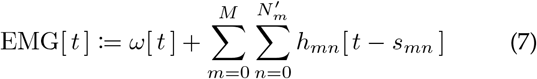

where we denote the m^th^ MU’s amount of MUAP observations using our revised formulation from Eq. 5 as 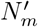. The n^th^ MUAP observation of the m^th^ MU is denoted as *h*_*mn*_[ *t* ] with it corresponding spike being *s*_*mn*_.

Finally, to extend our EMG signal to multiple electrodes, we also need to adjust our MUAP observation model so that it is now a vector of sequences as defined in Eq. 6. The length of the vector depends on the number of electrodes used and can be written as follows

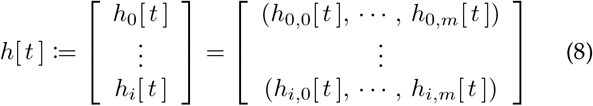

With this adaptation we can write the EMG signal of the i^th^ electrode as

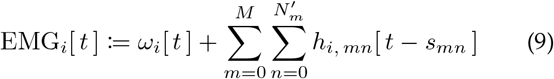

where *ω*_*i*_[ *t* ] is the additive noise for the i^th^ electrode and *h*_*i, mn*_[ *t* ] the n^th^ MUAP observation of the m^th^ MU picked up by the i^th^ electrode.

Now that we have an EMG signal formulation (Eq. 9) that models multiple electrodes, multiple MUs, and most importantly, multiple MUAP observations per MU, we can define an EMG bin consisting of only one electrode grid as follows

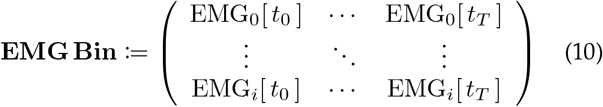

. In this definition we have *i* electrodes and the signals have been sampled at *T* different time points *t*.

#### 5.3.2 Explaining the deep learning choice

To extract information from an input signal that obeys Eq. 10, we need to be able to consider *J*| *J ≤ i* electrodes, 𝒯 | 𝒯 ≤*T* time points, and since we allow multiple MUAP observations to be present at the same time points and overlap each other, we must also look for the same pattern over the entire bin. So what we are interested in are specific MUAP overlap patterns that allow us to identify which movement will be performed. We assume that there are overlaps, i.e. several MUAPs that are summed up at the same time, as it is unlikely that only one MU is responsible for a movement in a healthy individual, especially if we assume that they perform their movements accurately.

The operation that can be used to extract these overlaps and at the same time meets our requirements is the same operation we used to construct our EMG signal model. The cross-correlation (adapted from Goodfellow et al. [57]) that we then need to use can be written for 2D as

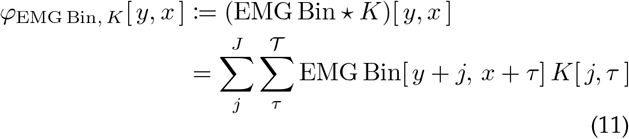

where *K* is the learnable kernel that covers *J* electrodes and 𝒯 time points.

If we then imagine that we have multiple learnable kernels, we can extract reoccurring MUAP overlap patterns over the length of the different movement recordings and use them to better understand the position of the hand. The neural network layers that use cross-correlation are called convolutional layers, and although they are called “convolutional”, most of them are implemented as crosscorrelation because this eliminates the need to mirror the kernels [57]. In the next part of this paper, we explain how our architecture is built and how we use these kernels to effectively and accurately extract kinematics from the sEMG.

### 5.4 Model

This work has been implemented in Python using PyTorch (version 1.11.0+cu113) [58] and PyTorch-Lightning (version 1.6.0) [59]. Fig. 9 shows a high level overview of the AI model architecture together with the input and output.

**Fig. 9:**
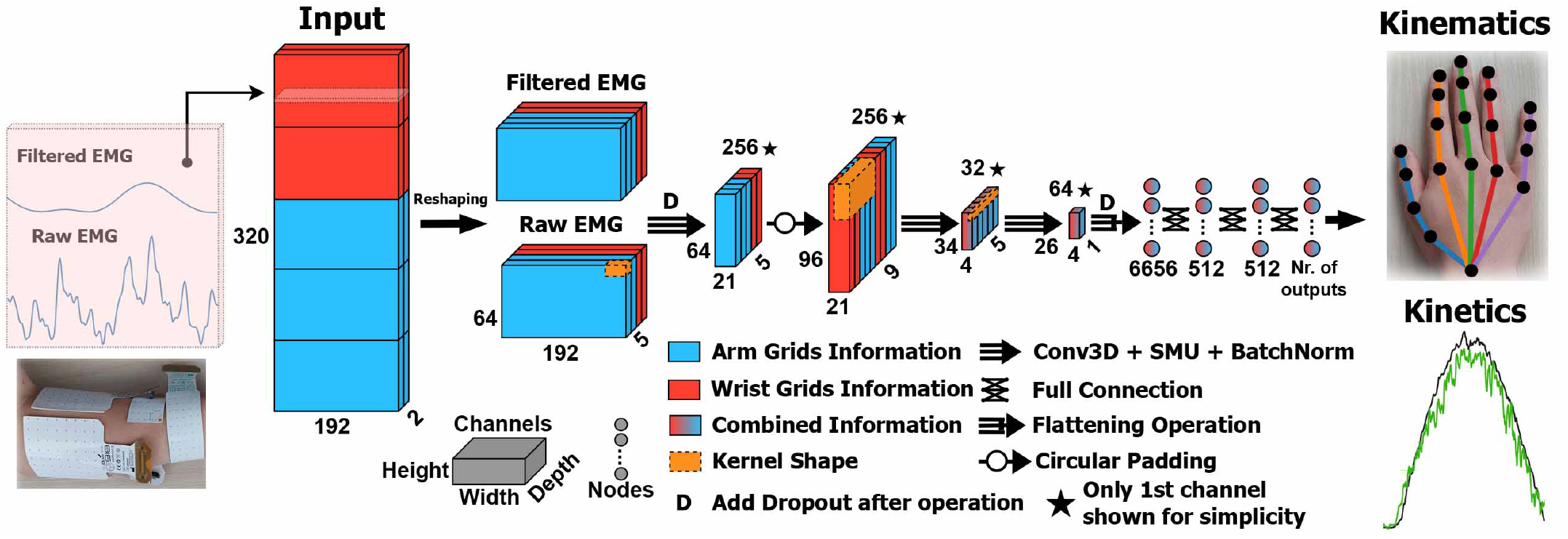
Architecture overview with a visualization of the circular padding.

#### 5.4.1 Architecture

In this paper, we use an architecture that is similar to the one we used in our previous conference paper [31]. We have already tested that architecture in a simulated real-time environment and found it to have satisfactorily inference latency.

The input is one or more sEMG matrices of shape 320 (total number of electrodes) *×*192 samples (*≈*94 ms). For this study we used two matrices. The first matrix contains band-passed (10 - 500 Hz) sEMG (“Raw EMG” in Fig. 9) and a second one that is a 20 Hz low-passed filtered copy of the first one (“Filtered EMG” in Fig. 9). These two matrices are concatenated in depth, giving us an input of shape 2 *×* 320 *×* 192. The input is then split by the amount of electrodes per grid (in our case 64 per grid) inside the model into a 4D tensor of shape 2 (number of inputted matrices) *×* 5 (number of grids) *×* 64 (number of electrodes) *×* 192 samples.

The output depends on the required format. In this study, we trained our network to output the entire hand skeleton in Euclidean space, all joint angles and the force applied to a force sensor.

When the whole hand has to be output in 3D, the output is a 60D vector (20 labels *×* 3 coordinates = 60 values). The wrist is stabilized and does not need to be output by the model as it has a constant position.

For the output of the angles, we have chosen to model the degrees of freedom of the hand as close as possible to the anatomical reality (Fig. 1C), not limiting ourselves to making logical mathematical simplifications when these have no effect on the hand model. We would therefore like to explicitly state that we are aware that anatomically the thumb consists of the carpometacarpal joint, the metacarpophalangeal joint and the interphalangeal joint (thumbs 3, 2 and 1 in Fig. 1B), but the overall position of the thumb can just as well be encoded by two 2D ball joints located at the position of thumb 2 and 1, as can be seen in Fig. 1C. With our hand model we get a 22D vector as output for the angle-based representation.

In the case of force prediction, only one variable needs to be output, which means we get a 1D scalar.

The model (Fig. 9) begins with a 3D convolutional layer (kernel size = 1x1x31; stride = 1x1x8; output channels = 256). We search each electrode in a window of 31 samples (*≈*15.2 ms) with a shift of 8 samples (*≈*4 ms) for action potentials in the sEMG.

Each 3D convolution used is followed by 3D batch normalization and the learnable activation function SMU [60], which has been shown to perform better than most common activation functions such as ReLU, Swish or Mish [60].

Before the next layer, we added a 3D dropout layer (p = 0.25) to prevent overfitting.

The next step is a circular padding. This layer extends a specified dimension by looping around to the other side of the same dimension. Fig. 9 illustrates the circular padding more clearly. The circular padding (2 for back and front; 16 for top and bottom) is applied to the output of the first step and ensures that the synchronicity of the recording is preserved for the next 3D convolution.

The subsequent layer is again a 3D convolution (kernel size = 5x32x18; dilatation = 1x2x1; output channels = 32). Dilation is used to look at events that are further apart. The kernel looks at five grids at once, giving us the grid combinations (1 is the first and 5 the last grid) of 45123, 51234, 12345, 23451 and 34512. This layer acts as an encoder and creates a representation of the extracted action potentials with lower dimensionality.

The last 3D convolution step’s (kernel size = 5x9x1; output channels = 64) task is to collect refined information and discard unimportant ones.

Afterwards, the output is flattened, has an 1D Dropout (p = 0.4) applied to it, and passed through a simple 3-layered perceptron (512, 512, number of output variables). The first two layers of the perceptron are followed by an SMU, while the last layer is followed by nothing, as this is the output.

We believe that the convolutional neural network learns the meaningful EMG information and represents it in high-dimensional space. The multilayer perceptron is then used to map the high-dimensional space into kinematics. Although we cannot definitively prove that this actually happens, in Fig. 7 we show results that strongly support the idea that the latent space contains information about the anatomy of the hand. This means that the multilayer perceptron no longer needs to learn what movement is being performed, but only needs to map it to the kinematics, which would support our intuition described above.

#### 5.4.2 Loss function

The loss function used is the mean absolute error (L1Loss in PyTorch). We chose this function because we want to linearly penalize any deviation from the ground truth.

#### 5.4.3 Optimiser and learning rate scheduler

For this architecture, we use the AdamW optimiser described in [61] with the AMSGrad correction from [62]. The weight decay is 0.01. We used the one cycle approach described in [63] as the learning rate scheduler with the upper bound being 10^−2.5^, the lower bound 10^−7^ and the initial learning rate 10^−4^.

### 5.5 Postprocessing

All output forms are filtered with a moving average filter of 150 ms. This is used to smoothen the output and eliminate the fast peaks in the output while preserving the overall trend.

For part of the 3D points analysis, we use the Procrustes method [64] shown in Algorithm 1.

#### Algorithm 1

Procrustes Analysis

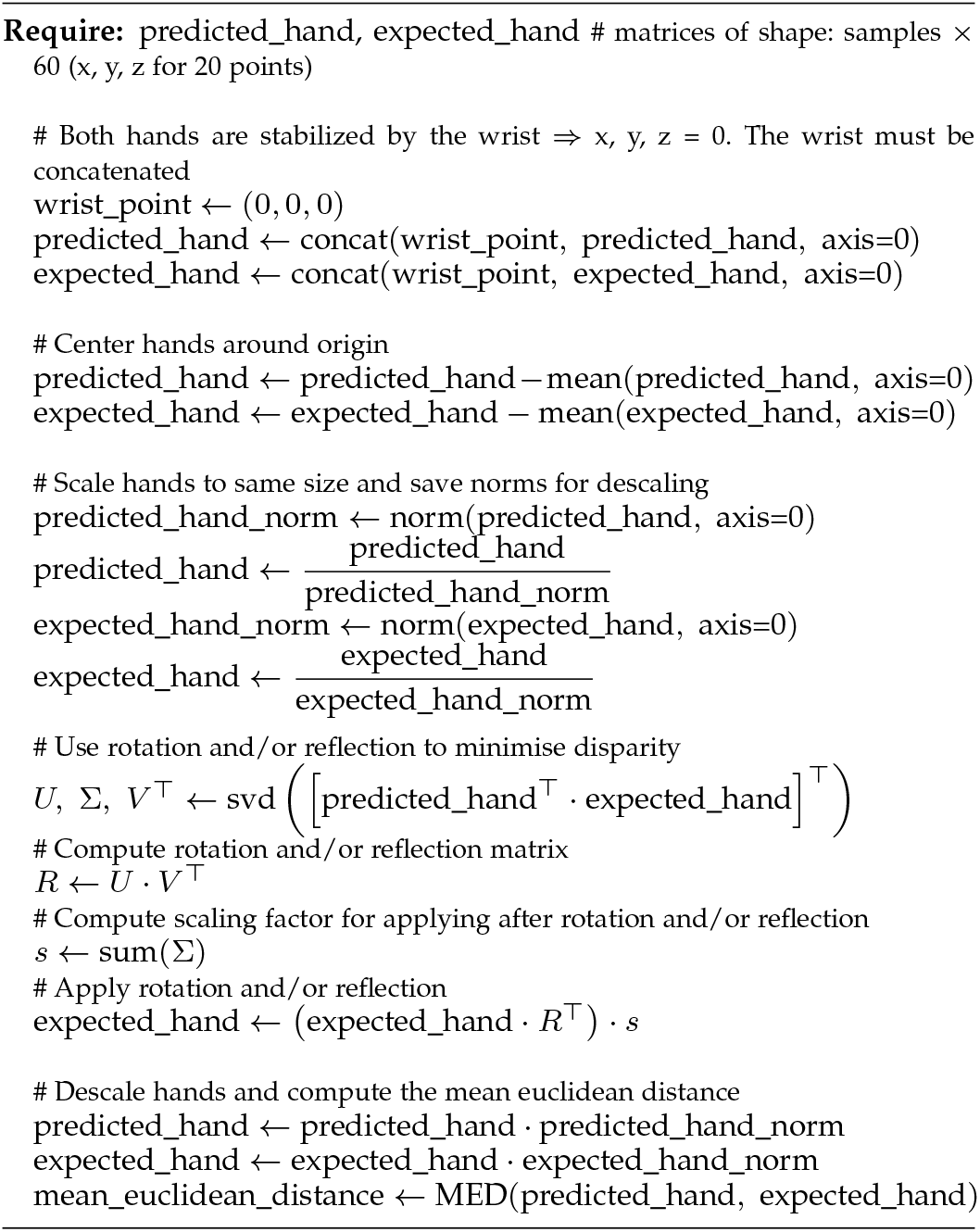

## 6 Data availability

All data generated or analysed during this study are included in the manuscript. Both the ground truth and prediction data used for making Fig. 2-6 is available at https:/doi.org/10.6084/m9.figshare.20481123.

The raw images (camera data as shown in Video 1-4) and monopolar EMG signals will be made available upon reasonable request.

The model for subject 1 can be tested online on a few movements from the test set and on the random movement set at https://huggingface.co/spaces/RaulS/D-Pose

## Acknowledgments

The authors gratefully acknowledge the scientific support and HPC resources provided by the Erlangen National High Performance Computing Center (NHR@FAU) of the Friedrich-Alexander-Universität Erlangen-Nü rnberg (FAU). The hardware is funded by the German Research Foundation (DFG).

Fig. 1A-C, 2A, 3A, 5A, 7A, and A1 are created with BioRender.com for which we wish to acknowledge it.

## Appendix A

### Supplementary Figures

**Fig. A1:**
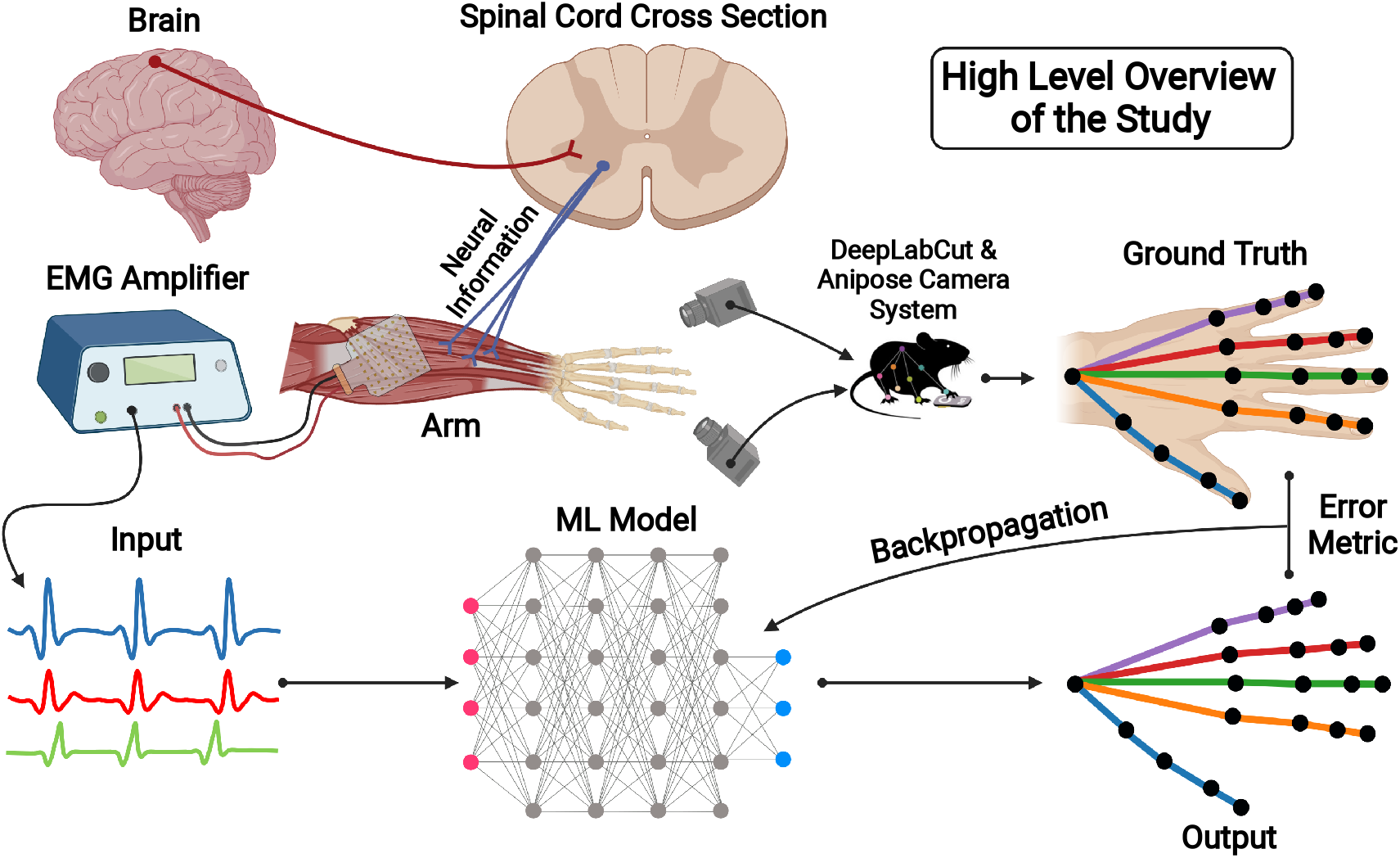
Extended high level overview of the study showing the neurophysiological pathway used for EMG generation (brain −→ spinal cord −→ arm muscles) and the framework used for the training of the ML model.

**Fig. A2:**
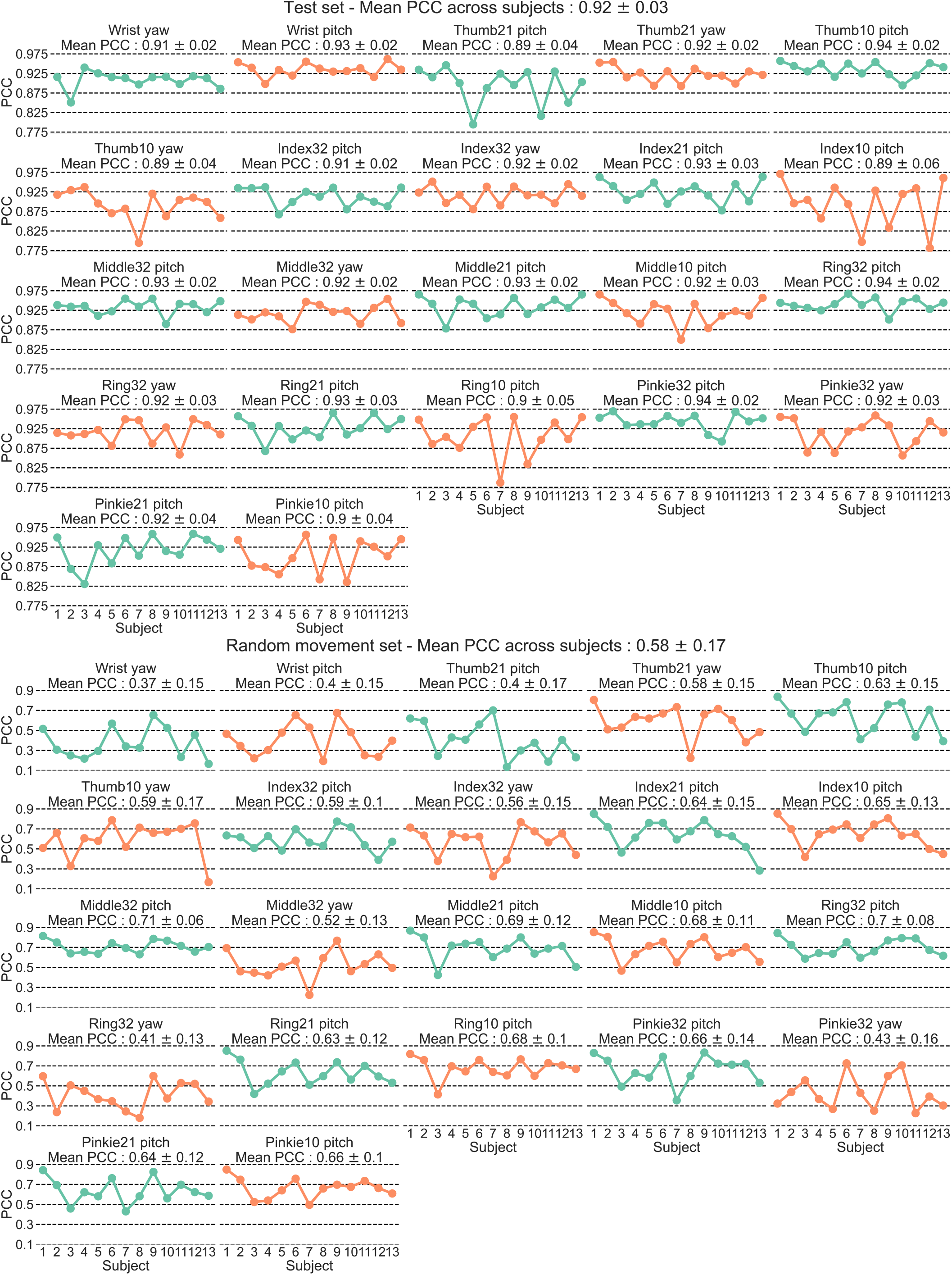
Individual joint angle correlations across subjects.

**Fig. A3:**
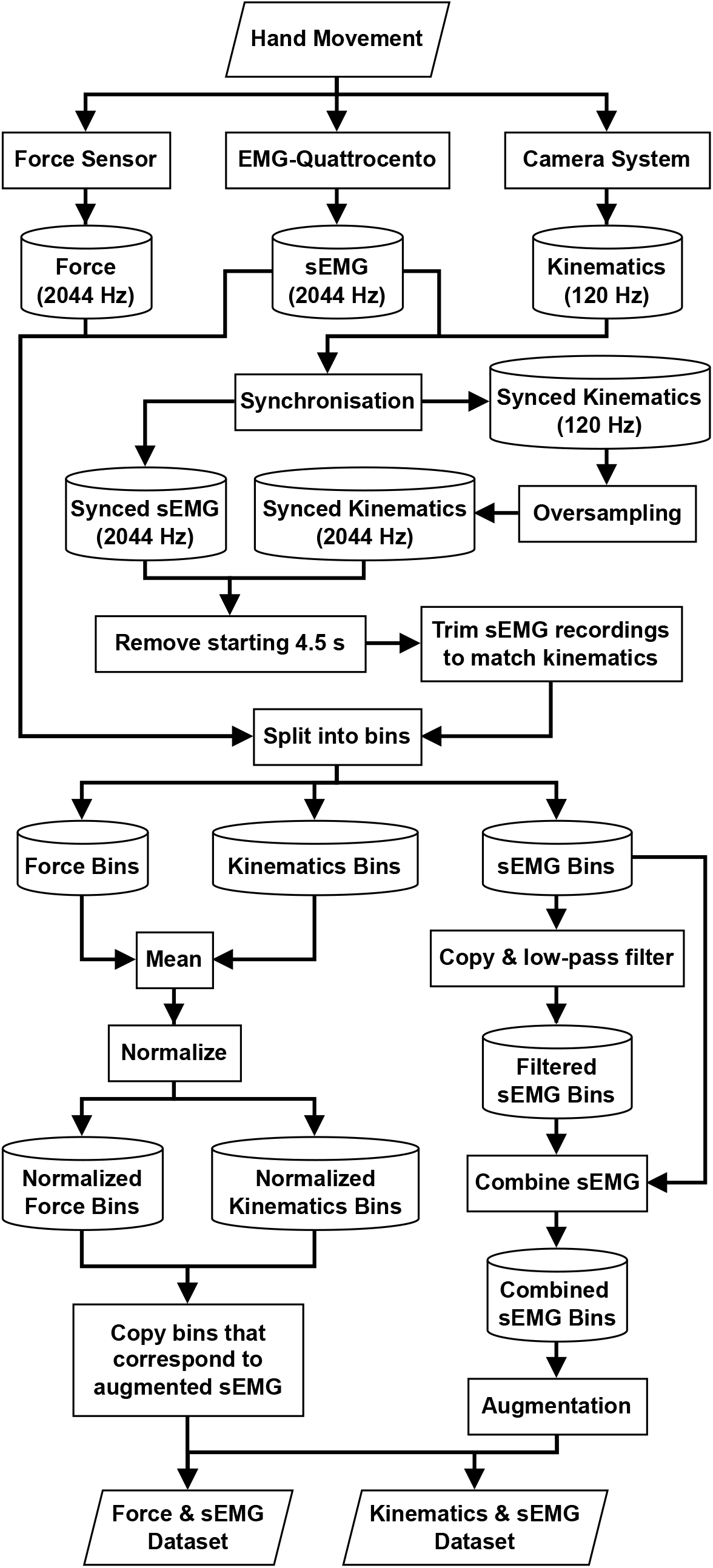
Data preprocessing flowchart.

**Fig. A4:**
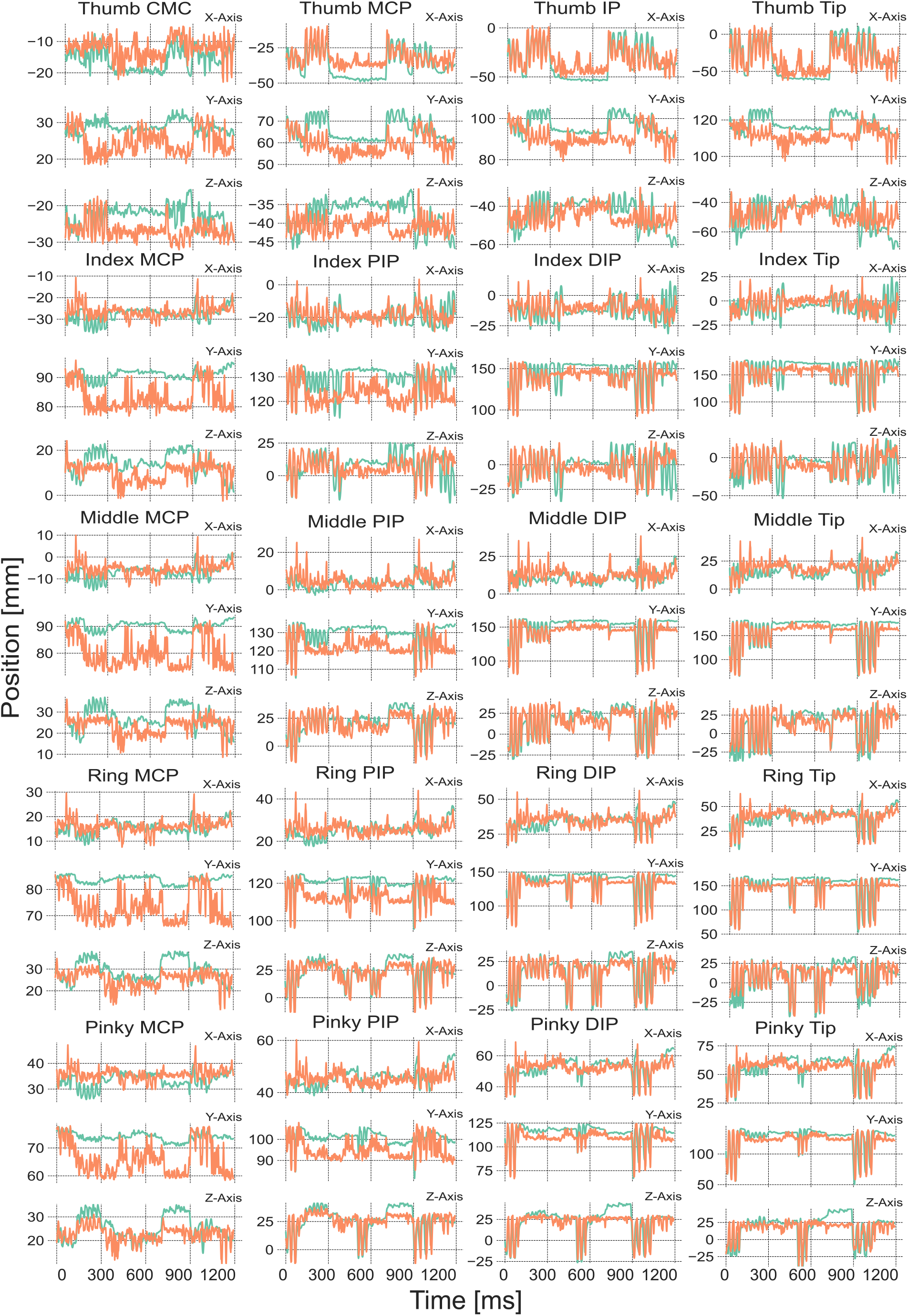
Comparison between expected and predicted joint positions from subject 10 on the random movement set without Procrustes analysis. Wrist position is not shown as the hand is stabilized by it and is set to be at (0, 0, 0).

**Fig. A5:**
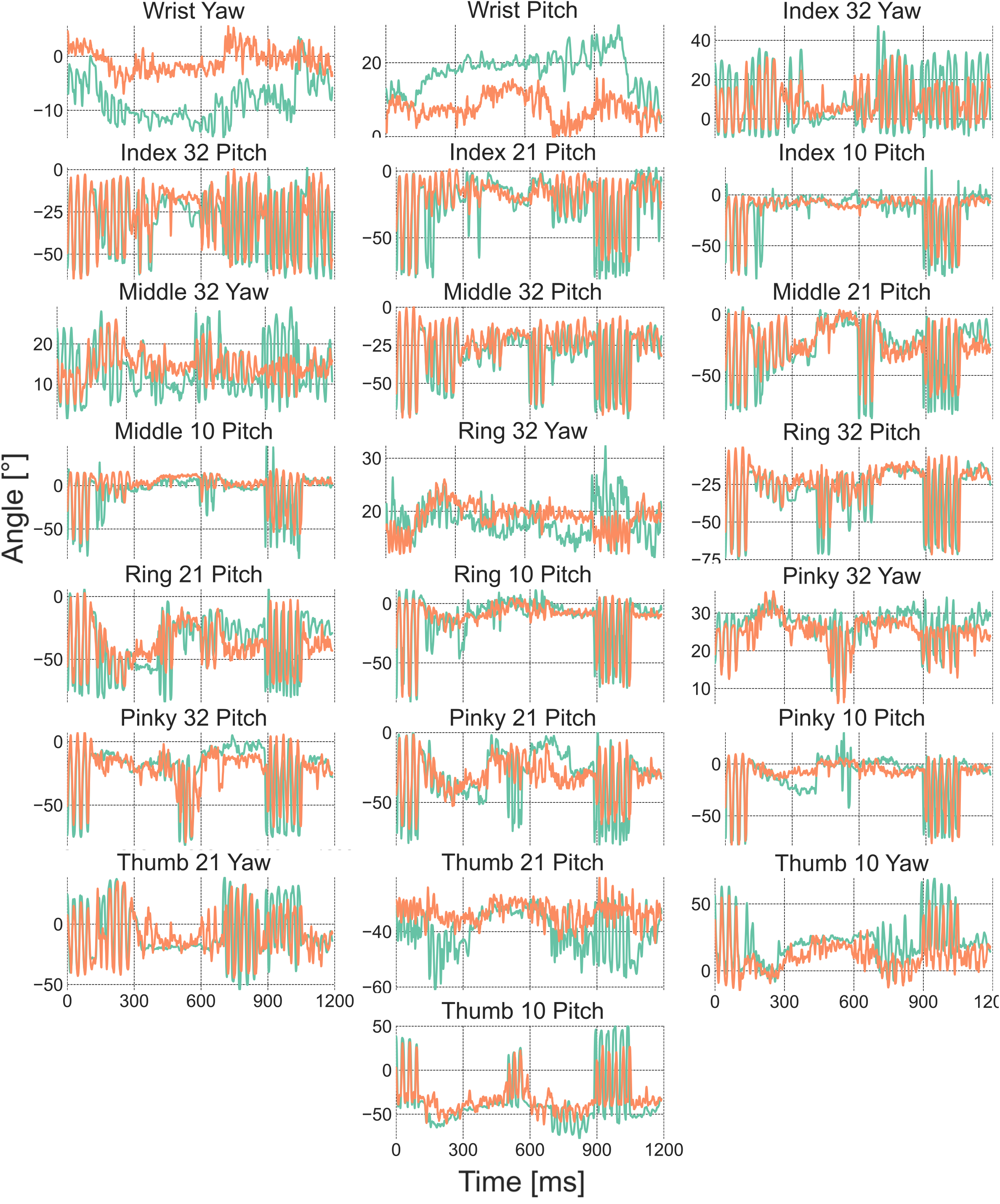
Comparison between expected and predicted joint angles from subject 10 on the random movement set.

